# Demonstrating Soft X-Ray Tomography in the lab for correlative cryogenic biological imaging using X-rays and light microscopy

**DOI:** 10.1101/2024.12.23.629889

**Authors:** Stephen O’Connor, David Rogers, Maryna Kobylynska, James Geraets, Katja Thaysen, Jacob Marcus Egebjerg, Madeleen C. Brink, Louisa Herbsleb, Michaela Salakova, Leon Fuchs, Frauke Alves, Claus Feldmann, Axel Ekman, Paul Sheridan, William Fyans, Tony McEnroe, Fergal O’Reily, Kenneth Fahy, Roland A. Fleck, Daniel Wüstner, Jeremy C. Simpson, Andreas Walter, Sergey Kapishnikov

**Author notes:** S.O’C. and D.R contributed equally to this work.

## Abstract

Soft X-ray tomography (SXT) enables native-contrast three-dimensional (3D) imaging of fully hydrated, cryogenically preserved biological samples, revealing ultrastructural details without the need for staining, embedding, or sectioning. Traditionally available only at synchrotron facilities, recent advances in laser-driven plasma sources have led to the development of compact soft X-ray microscopes, such as the SXT-100. The SXT-100 achieves imaging resolutions down to 54 nm full-pitch, with tomograms acquired in 30 minutes to two hours. Integrated with an epifluorescence microscope, the SXT-100 facilitates correlative workflows by bridging fluorescence and electron microscopy while preserving the structural integrity of vitrified samples. We demonstrate the capabilities of the SXT-100 through various use cases, including imaging *Euglena gracilis, Saccharomyces cerevisiae* yeast cells, and nanoparticles in mammalian cells. The relatively short tomogram acquisition times, the virtually non-destructive nature of soft X-ray tomography, and its quantitative imaging capabilities underscore its potential as a powerful tool for advanced biological imaging. Future developments promise enhanced throughput and deeper integration with emerging correlative imaging modalities, and a wider variety of sample types including tissue.

## Introduction

In this article, we introduce a lab-based soft X-ray tomography microscope (Fig. 1) utilising a high-brightness laser-driven plasma source with a metal target. Representing a major breakthrough in lab-based X-ray bioimaging, this microscope is being demonstrated in various applications, including virology, immunology, cell biology, and nanomedicine. Its versatility, compact design, and the non-destructive nature of soft X-ray tomography also facilitate the development of correlative workflows involving light and electron microscopy.^16,17^ For example, large field of view 3D volumetric data can add structural context to cryo-EM, while also offering a native unbiased view of the cell. Until recently, cryogenic soft X-ray tomography was available exclusively at synchrotrons (XM-2, ALS; MISTRAL, ALBA; B24, Diamond; U41-TXM, BESSY-II; BL08U1-A, SSRF; and 24A, NSRRC) due to the high brightness of X-rays required for imaging. Despite this limited accessibility, the synchrotron-based SXT user base continues to grow, with the technique validated in a growing number of applications, spanning chromatin rearrangement, virus–host interactions, cell motility, parasite life cycle, lymphocyte activation and function, among others.^9^ However, advancements in laser-driven plasma sources have led to the development of lab-based microscopes capable of performing soft X-ray tomography. Previous iterations using nitrogen gas puff or liquid nitrogen jet as the target for laser-driven plasma generation produced high-quality tomograms over reasonable exposure times and represent another approach that, due to a combination of engineering and fundamentals challenges, has not been developed into a commercial product.^10–12^

**Fig 1.**
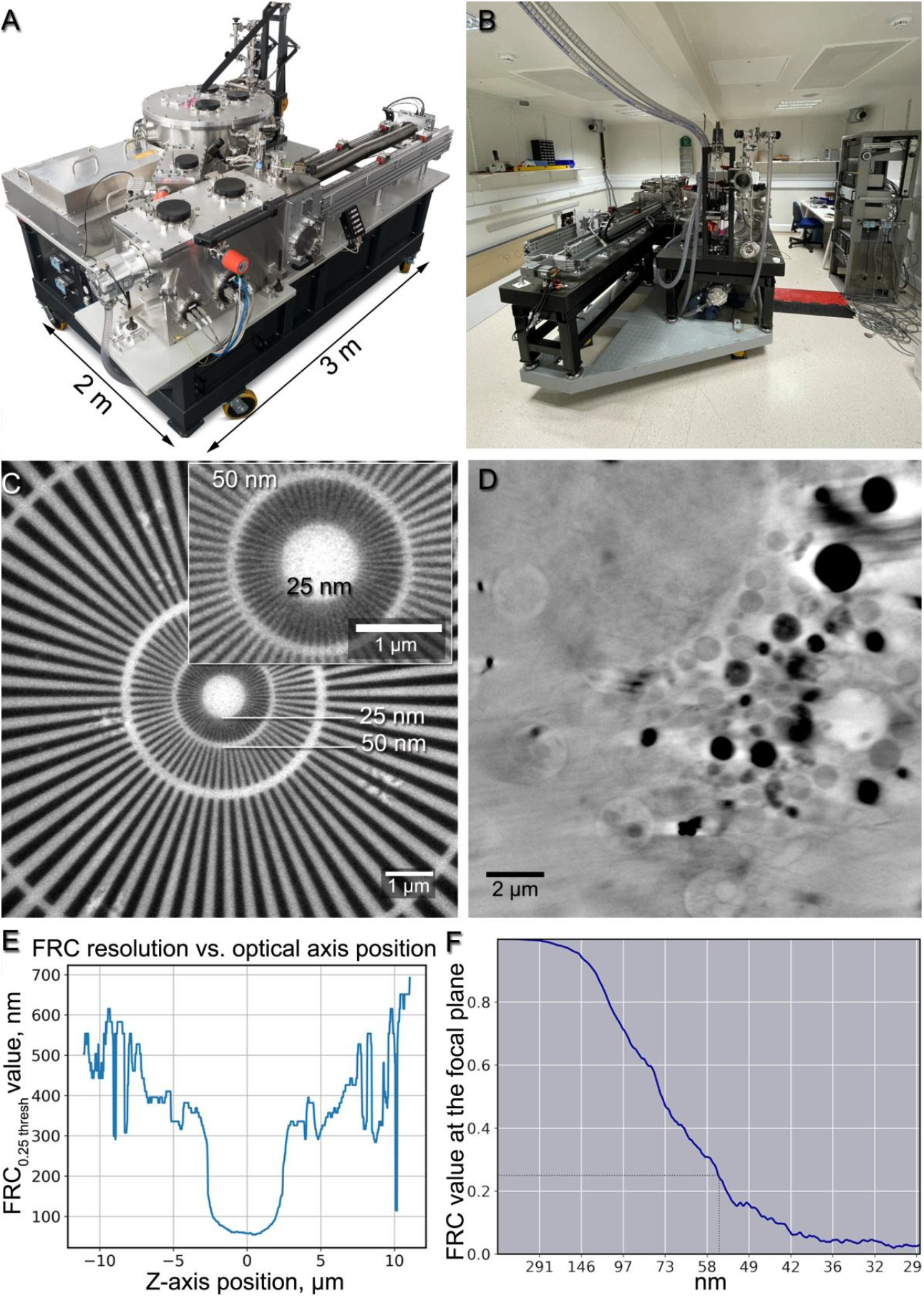
A lab-based soft X-ray microscope, Siemens star and Fourier-ring correlation resolution measurements. (A) The commercially available lab-based soft X-ray microscope, the SXT-100, developed by SiriusXT. (B) The SXT-100 in the Conway Institute for Biomolecular and Biomedical Research Imaging Core facility, University College Dublin. (C) A 50-second exposure soft X-ray projection through a Siemens star with the inner ring lines of 25 nm. (D) A focal-plane slice from a 3D reconstruction of a mammalian cell tomogram, acquired by the SXT-100 with a 1.5-hour exposure time over ±50° tilts, with a 1° tilt step. (E) FRC resolution measured at various distances from the focal plane in the 3D reconstruction. (F) FRC measurement of the focal plane slice indicating ca. 55 nm resolution.

In the following sections, we present soft X-ray tomograms, accompanied by correlative light fluorescence images, of various biological cells routinely acquired for research projects including viral disease research, lipid metabolism, nanoparticle delivery, and others.

## Results

### Lab-based soft X-ray microscope

Soft X-ray tomography (SXT) is a non-destructive three-dimensional (3D) imaging method that provides native-contrast visualisation of fully hydrated, cryogenically preserved biological samples. It reveals ultrastructural details without the need for staining, embedding, or sectioning. This imaging approach makes use of the unique absorption properties of X-rays within the “water-window” energy range (280-530 eV), where carbon-rich structures such as lipid layers and proteins strongly absorb X-rays while water absorbs them weakly. These properties allow for high-contrast imaging of biological specimens with the spatial resolutions down to lower tens of nanometres as reported in Oton et al.^1^ The technique is equivalent to medical Computed Tomography (CT) performed at the cellular level. Similar to Hounsfield units in medical CT, the data can be quantified based on soft X-ray absorption coefficients.^2,3^

The lab-based microscope, the SXT-100, developed by SiriusXT (see Fig. 1), achieves 3D resolutions in biological cells down to 54 nm full-pitch, while resolving Siemens star lines and spaces as fine as 25 nm, and enables the efficient acquisition of synchrotron-quality soft X-ray tomograms of fully vitrified cryogenic samples. This lab-based soft X-ray microscope features a footprint of 2×3 metres and includes an integrated cryo-epifluorescence microscope. Typical acquisition times for a single-cell tomogram range from 30 minutes to two hours, depending on sample thickness. In comparison to other volume imaging techniques that assess the ultrastructure of single cells, such as focused ion beam scanning electron microscopy (FIB-SEM), this allows for high-throughput data for quantitative statistical analysis. While a typical FIB-SEM dataset can resolve structures up to 5 nm, the acquisition of a single cell at such a resolution can easily require 48 hours, after which the sample is completely destroyed. Soft X-ray tomography, on the other hand enables subsequent, targeted high resolution imaging with, e.g., transmission electron microscopy (TEM) or FIB-SEM at cryogenic or room temperatures. The lab-based microscope accepts flat specimen holders, such as standard electron microscopy grids, and cylindrical specimens, such as thin-walled glass capillaries^15^ or cryo-FIB-SEM milled tissue pillars, for full-tilt tomography. Typical tilt series acquired in 1-degree steps over a 120-degree sample tilt range for flat specimens or 180-degree tilts for cylindrical specimens, collecting in total 121 or 181 projections, respectively. Each tilt exposure time varies from 15 to 60 seconds. An automated stack alignment routine performs fast and fiducial-free alignment before the 3D volume is reconstructed using standard approaches, e.g. weighted back projection (WBP) or Simultaneous Iterative Reconstruction Technique (SIRT) implemented in tomo3D software.^16,17^

To achieve high-resolution imaging of hydrated cells, cryogenic vitrification is necessary.^4,5^ This process involves rapidly freezing cells using techniques commonly employed in cryo-electron microscopy, such as plunging cells into liquid ethane cooled by liquid nitrogen.^6–8^ No cryoprotectants are required, simplifying sample preparation. However, it is crucial to maintain cells at cryogenic temperatures after freezing to preserve their structure and prevent ice crystallisation or thawing. Cryogenic sample environment has therefore been fully implemented in the SXT-100.^9,10^

### Resolution measurements

The spatial resolution achieved by the lab-based soft X-ray microscope was evaluated through two distinct methodologies: direct imaging of a Siemens star featuring 25 nm fine structure and Fourier-ring correlation analysis of a tomographic reconstruction of a Huh7 hepatocyte-derived carcinoma cell.

A Siemens star is a standard optical resolution test target, featuring a pattern of spoke-like structures that decrease in size towards the centre of the target. In the Siemens star imaging, lines as small as 25 nm in width were successfully resolved, as shown in Fig. 1. Notably, the specific Siemens star used in this study exhibited an uneven duty ratio at its 50 - 25 nm ring (SI Fig. 1).

For Fourier ring correlation analysis (FRC),^20^ we conducted tomographic scans of a Huh7 cell employing multiple frames per tilt. Subsequently, the frames were divided into two noise-independent datasets following the Tomogram Acquisition scheme outlined in the Methods section. FRC measurements were then conducted for series of reconstruction slices cut parallel to the detector plane, as illustrated in Fig. 1 D–F. The FRC threshold value of 0.25 was selected to be consistent with a previously published optical characterisation of the ALBA synchrotron-based soft X-ray microscope at the MISTRAL beamline.^1^

Utilising a threshold set at 0.25 of the correlation values, roughly corresponding to half the signal-to-noise ratio of the full dataset and chosen to maintain consistency with previously published measurements for a synchrotron-based SXT (as detailed in Oton et al. study),^1^ we attained a full-pitch resolution of 54 nm at the focal plane, as depicted in Fig. 1 E,F. Furthermore, the depth of field was estimated to be approximately 4.5-5 micrometres, as shown in Fig. 1 E.

### Correlative imaging workflow and freely available imaging routine for the quick co-registration of SXT and epifluorescent datasets of the same cell

The spatial resolution of soft X-ray imaging makes it highly suitable for correlative workflows, bridging the resolution gap between fluorescence and electron microscopy, while providing a native-contrast 3D structural context for the studied regions of interest, without causing damage to the sample.

The typical workflow^21^ for grid screening and tomogram acquisition starts with a mosaic scan of the grid using an integrated epi-fluorescence microscope. This step provides a visible light overview of the grid, overlaid with fluorescence signals, allowing for a quick assessment of grid quality and cell locations. Next, low-magnification X-ray transmission images, either single or in a mosaic, are acquired to identify regions of interest (ROI) for subsequent tomography acquisition.

Additionally, low-magnification X-ray projection mosaic is used to find regions of interest for imaging with volume EM, cryo electron tomography or other imaging techniques. A 2D grid mosaic of a 1mm^2^ area can be achieved in 20 mins. The acquisition times for tomograms typically range from 30 to 120 minutes. 2D projections alone, acquired over 3-15 seconds, can typically deliver the necessary information for targeting cells and cell areas for EM imaging.

We have furthermore established a workflow that allows for the automated co-registration of the correlative imaging datasets for a single cell, acquired using both SXT and the epifluorescence microscope. The workflow is based on lipid droplets that are easily detectable in both fluorescence and SXT imaging channels and does not require the addition of fiducial markers: For fluorescence microscopy, lipid droplets are stained with BODIPY 493/503, while in SXT, the contrast arises due to their carbon-rich content. Following the automatic detection and segmentation of lipid droplets in both channels based on Gaussian and contrast limited adaptive histogram equalisation (CLAHE) filtering, thresholding and morphological filtering, a phase cross correlation is calculated as a first, coarse alignment step between the two modalities.^22^ This is followed by an iterative optimisation based on a cost function and the final co-registration based on a Radial Basis Function (RBF) (see Materials and Methods and Fig. 2). The accuracy of the workflow is evaluated using the leave-one-out approach as published previously.^23,24^ Using lipid droplets as natural fiducial markers allows for an accuracy of the co-registration that is well within the pixel size of the low-resolution modality (for the wide field fluorescence microscope of about 500 nm) since the lipid droplet radii range from a few hundred nanometres up to a micron. Furthermore, using lipid droplets instead of artificially added fiducial markers, comes with the advantage that these subcellular structures are endogenous to the cells and distributed throughout them.

**Fig 2.**
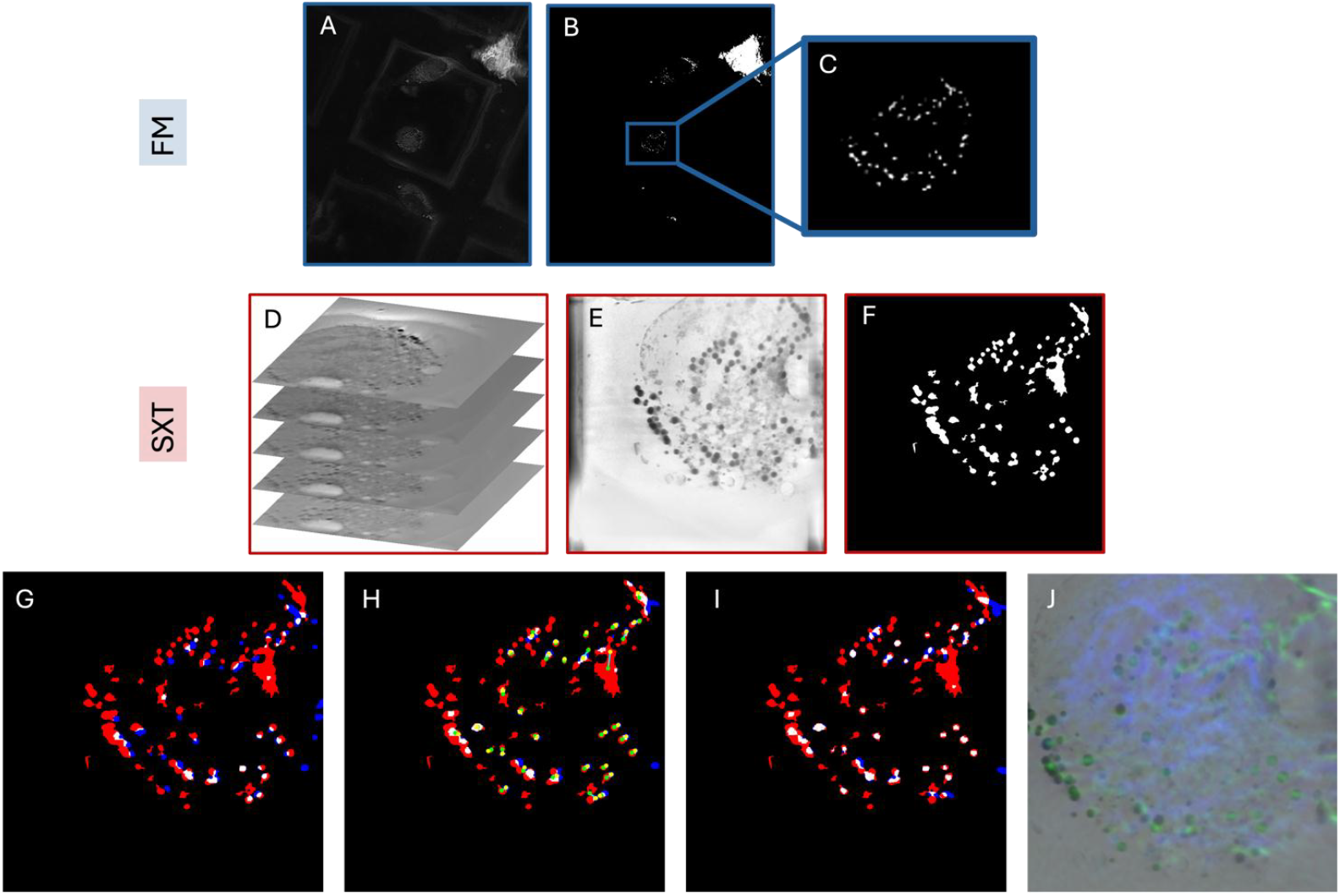
Illustration of the co-registration workflow without the addition of fiducial markers, based on endogenous lipid droplets, visible in both the SXT and wide field fluorescence channel (BODIPY staining). The red channel represents the X-ray image, the blue channel corresponds to the fluorescence image. Areas of overlap between the two modalities are displayed in white, indicating regions where both signals coincide. (A) Gaussian- and CLAHE-filtered wide field FM image. (B) Segmented lipid droplets of the FM modality based on global Otsu’s thresholding. (C) Selected Region of Interest to co-register with the SXT dataset. (D) Original SXT image stack with the unprocessed data. (E) 2D-projected and filtered SXT stack. (F) Threshold and size-based segmented lipid droplets. (G) The Phase Cross-Correlation (PCC) provides a first, coarse alignment between the X-ray (red) and fluorescence (blue) images. This step serves as the starting point for the alignment process. (H) Iterative Alignment with a Cost Function. Yellow points represent the centroids of the red segments, showing their calculated centers of masses. Each yellow point is connected to the corresponding blue segment that it aligns with, demonstrating the progress of the alignment process and the matched pairs between the two modalities. (I) The Final Warped Image is achieved using Radial Basis Function (RBF) interpolation. This method addresses local distortions to enhance the correspondence between the X-ray and fluorescence images, resulting in a more precise overlap of the two modalities. (J) Overlaid and co-registered minimum intensity projection of the SXT stack and the wide field FM image.

### Soft X-ray and cryo-light fluorescence Imaging of *Euglena gracilis*

A correlative light fluorescence and X-ray images of *Euglena gracilis* cells plunge frozen on a TEM grid are shown in Fig. 3. Euglena is a complex eukaryotic cell that requires effective ROI targeting of structures for higher resolution cryo-vEM and/or cryo-ET. Correlating ultrastructural information from the entire cell in FIB-SEM with the SXT reconstruction enhances our understanding of the contrast and appearance of organelles and structures in soft X-ray tomograms. Furthermore, the soft X-ray tomograms are highly informative for correlative studies because the volume is unaffected by ice, which can limit light and electron microscopies. In the soft X-ray tomograms shown in Fig. 3, absorption based contrast clearly discriminated major organelles. Clearly present were dorsal flagella (arrows), the combined structures containing the feeding apparatus and eyespot, including the ventral flagellum, ventral root, paraxial swelling and flagella pocket vestibulum (white open circle). Integrated fluorescence imaging was able to localise fluorescence associated with photosynthetic systems (chlorophyll fluorescence emission). When combined with the SXT volumes, chloroplasts with thylakoid structures (white squares) and non-fluorescent paramylon granules (white triangles) could be readily identified. Additionally, the nucleus could be clearly located in SXT images (white circle). This information can be employed to target sub cellular and potentially sub-organelle regions of interest for further higher resolution study by EM techniques.

**Fig 3.**
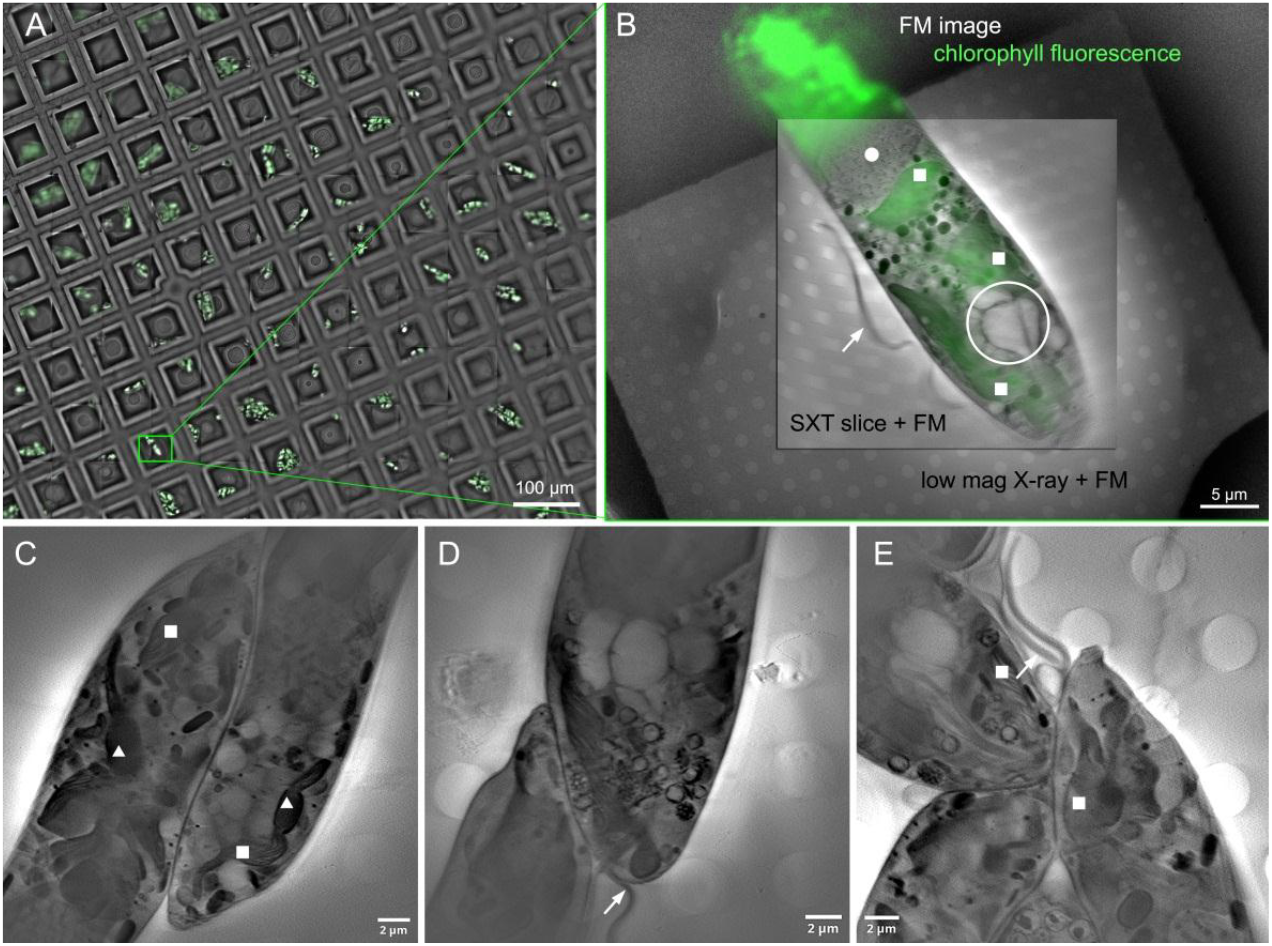
*Euglena gracilis* cells cryo-vitrified on 3 mm Quantifoil-coated TEM grids and imaged by cryo-SXT and the integrated fluorescence microscope in the SiriusXT table-top SXT microscope. (A) A visible light fluorescence mosaic of a portion of a 3 mm TEM grid showing grid windows and the chlorophyll fluorescence signal. The area where the soft X-ray tomogram was collected, as shown in panel B, is enclosed in a green rectangle. (B) A composite fluorescence microscopy (FM) image of the *Euglena* chloroplast overlaid with a low magnification X-ray projection image acquired with 10 s exposure and a virtual slice through a high magnification SXT tomogram acquired over 1.5 hours. (C, D, E) Virtual slices through tomograms with 29×29 μm^2^ field of view featuring seven *Euglena* cells.

### Soft X-ray tomography of cryo-vitrified yeast cells

The low magnification X-ray images reveal *Saccharomyces cerevisiae* yeast cells (Fig. 4) as distinct dark elongated objects, approximately 5 micrometres in size, set against the backdrop of the recognizable pattern of Quantifoil film holes. An example of a combined X-ray and light fluorescence dataset is presented in Fig. 4. A cropped section of an FM mosaic containing a few grid areas where cells were located is shown in Fig. 4A. A specific grid area with visible cells was chosen (green rectangle) and an FM image was acquired (Fig. 4B) followed by a mosaic of low magnification X-ray projections (Fig. 4C) or single low magnification X-ray projections (Fig. 4D) to identify ROIs for a tomogram acquisition. One such ROI is marked with a blue rectangle (Fig. 4D). A zero-degree tilt high magnification projection overlaid with a MSSR-processed^25^ FM image is shown in Fig. 4E, and a virtual slice, extracted from the three-dimensional (3D) reconstructed tomogram with the voxel size of 29 nm is shown in Fig. 4F.

**Fig 4.**
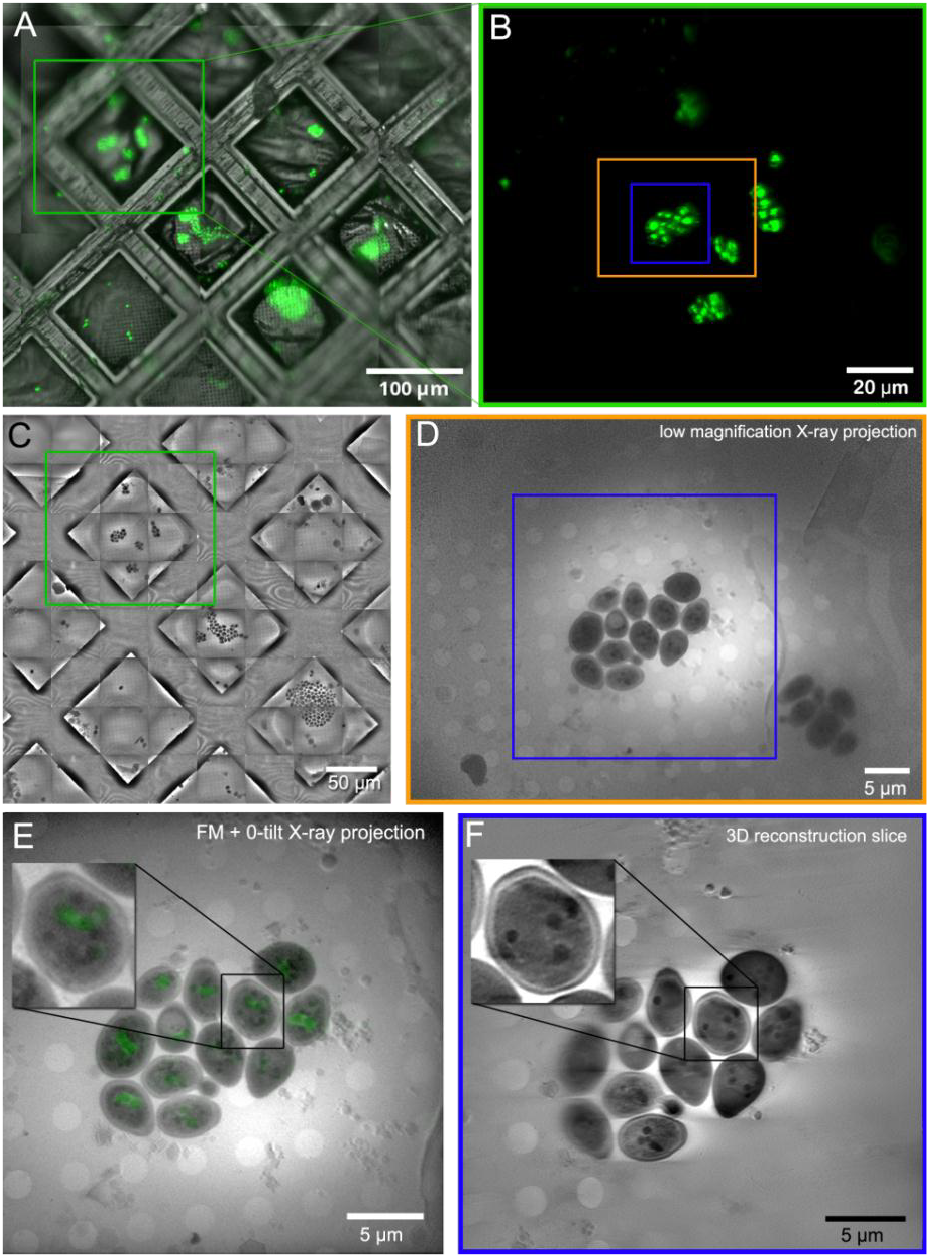
*Saccharomyces cerevisiae* yeast cells imaged using the SXT-100. (A) A portion of a visible light fluorescence microscope images mosaic of a 3 mm TEM grid with vitrified yeast cells labelled with BODIPY493/503 dye. (B) a single fluorescence image of a grid area containing a number of yeast cells. (C) A mosaic of low magnification X-ray transmission images covering an area of 370 × 350 μm^2^ acquired in less than 5 minutes. (D) A single low magnification X-ray transmission image capturing yeast cells within a 43×58 μm^2^ field of view on a grid window acquired with a 10-second exposure. (E) Zero-tilt X-ray tomogram projection displaying a 30×30 μm field of view overlaid with a MSSR-processed fluorescence image. (F) A virtual slice extracted from the reconstructed tomogram acquired over one hour.

Within yeast cells, discernible submicron-sized dark spherical objects, visible both in X-ray transmission images and the reconstruction were observed. These correspond to lipid bodies, demonstrating pronounced X-ray absorption due to their high carbon content. By contrast, the least absorbing features within yeast cells were vacuoles, primarily containing digestive enzymes in solution. The ability of the SXT-100 microscope to change magnification and the field of view size not only facilitates screening of the sample (Fig. 4A-C), but also allows high-quality correlation of the necessary lower resolution of epifluorescence with X-ray images to be achieved (Fig. 4E). The 3D reconstruction of a high-resolution tomogram (Fig. 5) revealed the spatial arrangement of the droplets relative to the vacuole, enabling study of the process of vacuolar ingestion of lipid bodies during starvation, namely lipophagy.^26^

**Fig 5.**
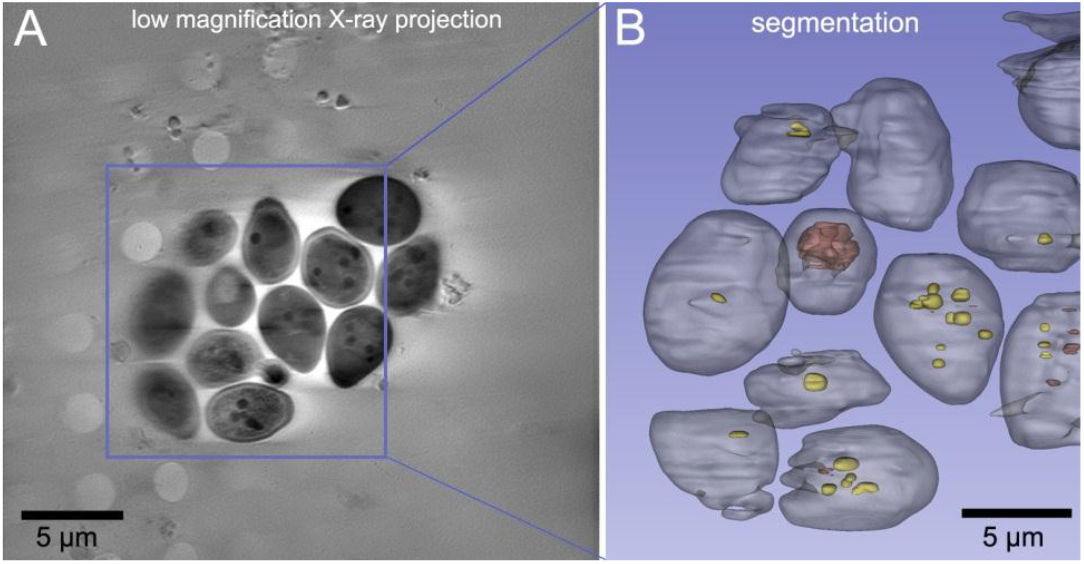
(A) A virtual slice extracted from the reconstructed tomogram featuring twelve *Saccharomyces cerevisiae* yeast cells. (B) Segmentation surface rendering of a cropped reconstruction region outlined with a blue rectangle. Yeast cells are rendered in grey, lipid droplets in yellow, and digestive vacuoles in brown.

### Imaging nanoparticles in cells using fluorescence and soft X-ray tomography

#### Polymeric nanoparticles in HeLa cells

Given the ability of the SXT-100 microscope to resolve subcellular structures in yeast cells, we next applied the system to a use case in cultured mammalian cells. Nanoparticles (NPs) are emerging as important therapeutic delivery vehicles, and gaining a greater understanding of their distribution in the subcellular environment is of high importance to improve their performance. As they are relatively well-characterised in the literature, 100 nm red Carboxylate-Modified FluoSpheres™ and HeLa cells (Fig. 6A) were used as a model. From previous work using fluorescence light microscopy approaches, these NPs are known to accumulate in lysosomes over time.^27^ HeLa cells were pulsed with NPs for 4 hours, incubated with Hoechst 33342 (to identify cell nuclei) and LysoTracker (to identify lysosomes), and plunge frozen for cryo-soft X-ray imaging. Red 100 nm FluoSpheres (Fig. 6 B1) were detected at the perimeter of the HeLa cell, this was confirmed by correlating the SXT with the FM image. This also showcased the ability of the lab-based soft X-ray microscope to resolve 100 nm particles. We observed that the FluoSpheres (red) accumulated in lysosomes (green) (Fig. 6 B2), which was consistent with previous studies.^27^ In Fig. 6 B2 and B3 mitochondria and lipid droplet morphology can be clearly discerned; this is made possible by exploiting the strong absorption of soft X-rays by carbon dense cellular moieties and is in agreement with that observed in transmission electron micrographs.^28^ Soft X-ray tomography enables cells to be imaged in their whole, native state, which allows the capture of rare events in the mesoscale to create a more complete picture of how NPs traverse cells when correlated with light microscopy. Understanding the intracellular trafficking of NPs is critical to ensure effective drug delivery when designing novel nanomedicines.^27,28^

**Fig 6.**
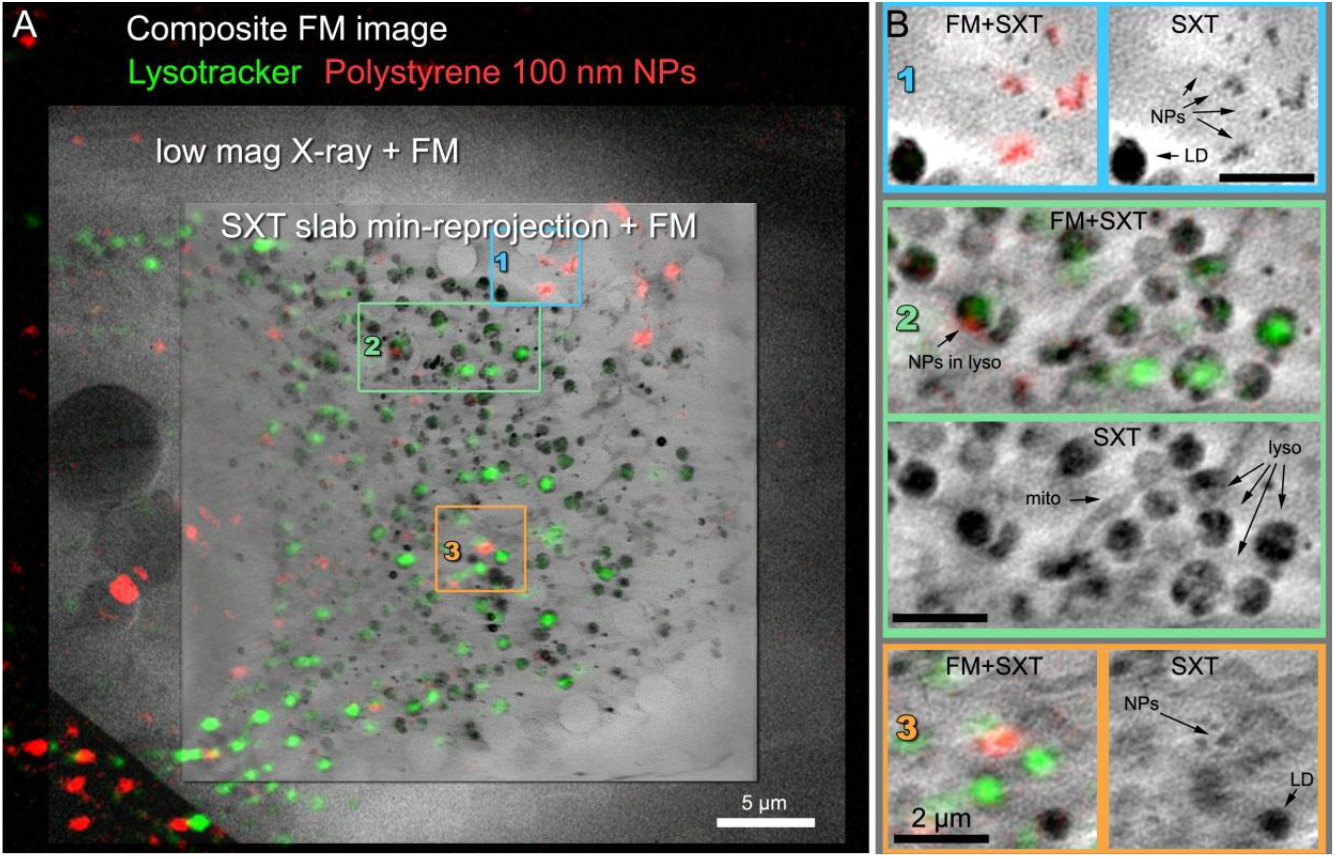
A HeLa cell incubated for four hours with 100 nm fluorescent polystyrene nanoparticles. (A) Composite fluorescence microscopy (FM) image of Lysotracker and fluorescent nanoparticles overlaid with a low magnification X-ray projection and a three high magnification SXT slices across the cell, (B,C) Zoom-in of selected areas in the cell cytoplasm showing the SXT slice (top of each panel) and the same SXT slice overlaid with the composite FM image (bottom of each panel).

#### Inorganic-organic hybrid nanoparticles for cancer therapy

Nanoparticles exist in a variety of sizes and materials for therapeutic delivery. Another example is zirconyl-containing inorganic-organic hybrid nanoparticles (IOH-NPs), previously described by Heck et al.^29^ These IOH-NPs consist of an organic drug/dye anion and an inorganic cation such as zirconyl ([ZrO]^2+^) that together result in nanoparticles being insoluble in water. Due to their metal-containing structure (and hence higher absorption of X-rays) and their incorporation of a red fluorophore (DUT647: Dyomics-647-uridine triphosphate), the IOH-NPs were found to be visible and detectable using both SXT and FM. With a diameter of ca. 60 nm, single IOH-NPs were resolvable using SXT. These nanoparticles have specific features and advantages of the fluorescent [ZrO]^2+^[(CMP)_0.99_(DUT647)_0.01_]^2-^]^2^ IOH-NPs^-^ (CMP: cytidine monophosphate), that were originally developed at the Karlsruhe Institute of Technology, are: *i*) synthesis in water, *ii*) high load of drug anion and/or fluorescent dye anion (up to 80 wt-% of total nanoparticle mass; here with CMP representing a potential drug anion and DUT647 as fluorescent dye anion), *iii*) high photostability, *iv*) intense emission, and *v*) flexible nanoparticle composition and fluorescence,. The therapeutic benefits of the high drug load of IOH-NPs in minimizing undesired side effects and overcoming mechanisms of chemoresistance have been demonstrated in a pancreatic cancer mouse model with gemcitabine monophosphate (instead of CMP).^30^ Due to fluorescence labelling, IOH-NPs facilitate monitoring *in vivo* and *ex vivo* IOH-NP distribution, confirming their accumulation within tumour tissue and their uptake in tumour cells via endocytosis, followed by intracellular trafficking to late endosomes/lysosomes.^30^ To understand the fundamental concepts of how the IOH-NPs interact with the biological systems at cellular and subcellular levels, how physicochemical properties influence intracellular uptake and trafficking and how this might affect nanodrug efficacy, there is a high need to explore the intracellular internalisation, transport of IOH-NP and drug release at a single-cell level. As shown in Fig. 7, this can be achieved by means of correlative fluorescence and soft X-ray-based imaging. To study the internalisation dynamics, murine triple-negative breast cancer cells (H8N8) ^31^ were incubated with the IOH-NPs and fixed at 4 hours after internalisation. SXT provided sufficient resolution to *(i)* assess the cellular organelles of each cell for each time point and characterise ultrastructural changes that were induced by the IOH-NPs, and to *(ii)* follow the IOH-NP uptake, fate, and co-localisation with organelles (such as lysosomes). Importantly, in contrast to higher-resolution imaging modalities, such as volume EM, SXT was able to provide throughput of 20-30 cell tomograms, and hundreds of low magnification projections per week, to visualise enough cells for each time point to quantitatively and statistically analyse the NP distribution and ultrastructural changes in a reasonable time frame of approximately 2 weeks.

**Fig 7.**
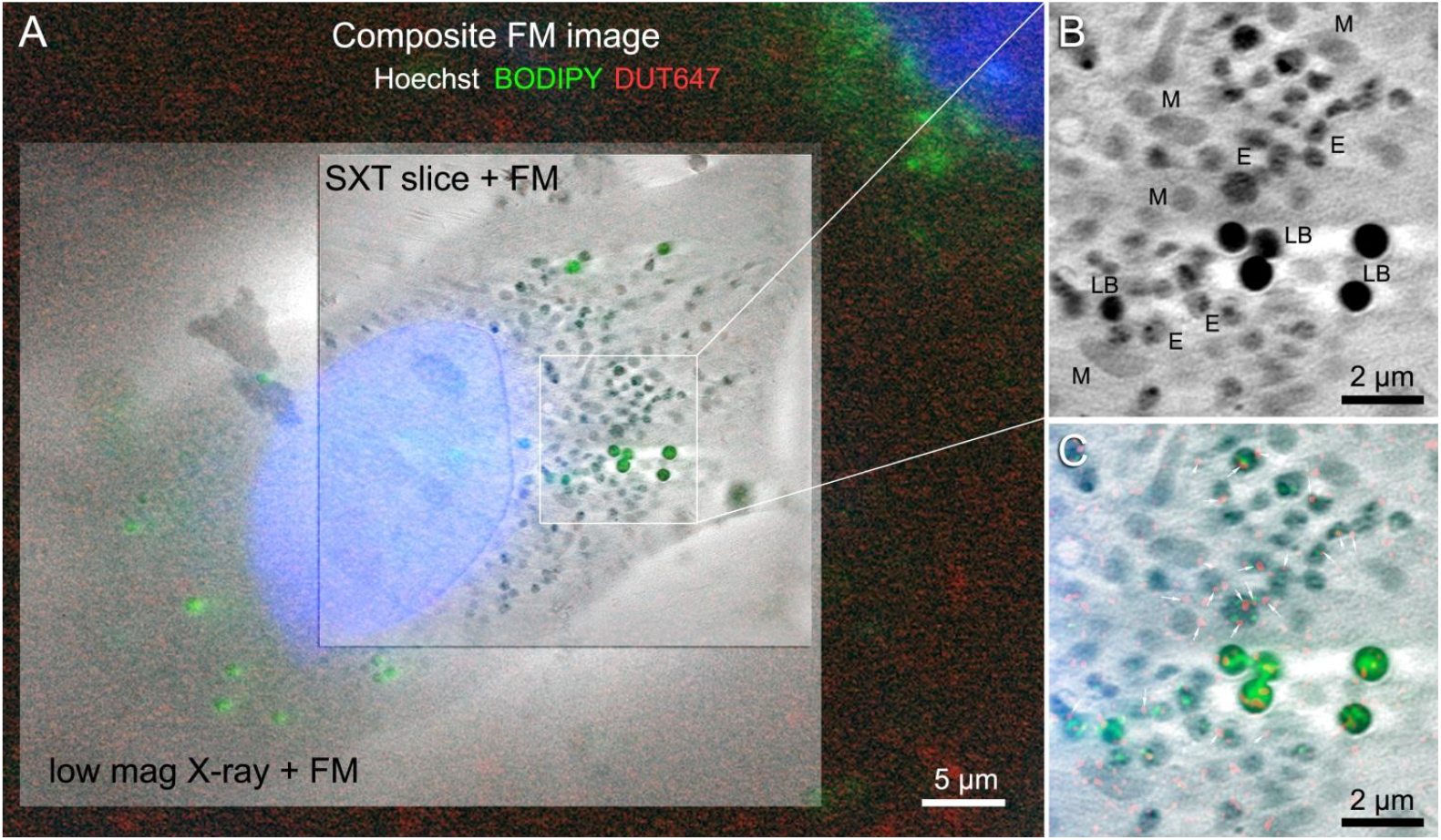
An H8N8 cell incubated for four hours with 50-60 nm [ZrO]^2+^[(CMP)_0.99_(DUT647)_0.01_]^2-^]^2^ IOH-NPs. (A) Composite fluorescence microscopy (FM) image of Hoechst 33342, BODIPY, and DUT647 overlaid with a low magnification X-ray projection acquired over 10 s and virtual slice through a high magnification soft X-ray tomogram acquired over 1.5 hours. (B) Zoom of a selected area in the cell cytoplasm showing the SXT slice. Abbreviations: LB - lipid bodies, M - mitochondria, E – endosomes. The appearance of most of the endosomes resembles that of lysosomes shown in Fig. 6. (C) Zoom of the same cytoplasm area showing the SXT slice overlaid with the composite FM image. White arrows point to the nanoparticles visible as red dots. The location of the nanoparticles coincides primarily with lysosomes, while a few nanoparticles co-locate with lipid bodies.

## Discussion

The development of a compact soft X-ray tomography microscope, which enables fast high-resolution imaging of fully hydrated, cryogenically preserved biological cells in a laboratory setting, represents a significant advancement in the field of soft X-ray tomography (SXT) and cellular biology. By making SXT accessible to a wide range of researchers at the local lab, core facility level, or commercially, independent of synchrotron facilities, this microscope opens new avenues for in-depth studies of cell structure across various research fields, as well as for correlative light, electron, and X-ray microscopy.

Our results demonstrate that the lab-based system achieves spatial resolutions comparable to those obtained at synchrotron facilities, specifically resolving structures as fine as 25 nm with the Siemens star and achieving a full-pitch resolution of 54 nm using Fourier-ring correlation analysis of reconstructed tomogram slices of biological samples. However, the lab-based SXT-100 typically requires 30 minutes to two hours to acquire a tomogram, compared to the 5-15 minutes required by synchrotron-based systems. On the other hand, the SXT-100 offers a larger field of view, up to approximately 45×60 μm^2^, due to its different illumination scheme. Furthermore, synchrotron-based systems can perform full-field absorption spectroscopy allowing for flexible changes in X-ray energy, while the laser-based source of the SXT-100 provides a few narrow peaks in the water-window X-ray energy range, the most intense being at 455 eV and 500 eV, pre-selected by the multilayer optics.

In addition to its high spatial resolution, the compact soft X-ray microscope offers several benefits for cellular and subcellular studies.

One of the primary advantages of SXT is its ability to provide native-contrast 3D structural imaging of whole cells in their close-to-native environment.^32^ The microscope’s capability to handle cryogenically vitrified samples at or below −160°C through all imaging and sample transfer steps prevents ice crystallisation,^6,9^ maintaining the integrity of vitrified biological samples. This enables visualisation of cellular organelles and ultrastructure without the need for staining or labelling. The 3D structural context provided by SXT is invaluable for correlating functional imaging data from fluorescence microscopy, as it allows for precise localisation and interpretation of fluorescent signals within the entire cellular architecture. An integrated epifluorescence microscope further enhances this capability by enabling fluorescence mapping of the specimen without removing the sample from the stage.

Furthermore, by offering direct 3D imaging of cell structures, integration of a lab-based soft X-ray tomography will significantly improve the throughput of EM and correlative light and electron microscopy (CLEM) by making it easier to locate structures of interest, thereby streamlining these workflows.

Given the ability of SXT to generate a 3D map of entire cells, the technique is better suited to capture rare events (e.g. nanoparticle endosomal escape), which might be missed when slicing through small portions of a cell by EM.^33^ The ability to visualise entire cells in 3D with native contrast allows researchers to identify and analyse these rare events within their full cellular context, which might otherwise be missed or overlooked using other imaging techniques. Importantly, within a week, almost 50 cells and different time points can be imaged. For instance, in our study of 100 nm nanoparticle-fed HeLa cells, we demonstrated the utility of this integrated system by combining soft X-ray tomograms with fluorescence images, allowing for precise localisation of nanoparticle markers within the cellular context. Similarly, in the study of [ZrO]^2+^[(CMP)_0.99_(DUT647)_0.01_]^2-^]^2^ IOH-NP distributions within H8N8 breast cancer cells, the ability to visualise the distribution and impact of nanoparticles on cellular morphology helps in understanding the intracellular uptake and transport mechanisms of nanoparticle-based drug delivery systems to treat cancer cells Dynamic studies over time will enable to assess not only the trafficking but also the degradation process of the IOH-NPs.

The ability of SXT to locate cells or specific areas under the sample ice, combined with providing a reliable estimate of ice thickness and quality, adds another layer of utility for sample preparation and quality control in cryo-EM studies. Given that the radiation dose from an SXT tomogram typically ranges from 4 to 100 MGy, equivalent to <1 to 20 cryo-ET projections, depending on chosen exposure and sample thickness, SXT offers a minimally invasive approach that complements downstream imaging.

Another important aspect of SXT is the quantitative nature of its absorption contrast, which follows the Beer-Lambert law.^2,4^ This enables the distinction of cellular structures or quantification of concentrations by measuring linear absorption coefficients from SXT reconstructions. This approach works particularly well for cylindrical sample geometries, such as thin glass capillaries loaded with cells,^34–36^ which are also supported by the lab-based microscope.^21^

Overall, the ability to resolve fine structures with high-throughput, combined with its native-contrast quantitative imaging capabilities^37,38^ and its role in facilitating complementary imaging techniques, underscores the versatility of compact soft X-ray tomography systems and strong potential for advancing research on cellular structure and function.

## Future Directions

The use of a compact laser-based metal target source to generate water-window soft X-rays in a full-field X-ray microscope will bring soft X-ray tomography into labs and core facilities by providing direct and instant access to this imaging modality. On-going developments in the lab-based SXT microscope, such as multiple grid handling, biosafety levels 2/3 sample shuttles, tissue slab imaging, and options for confocal, super-resolution fluorescence, and Raman spectroscopy, as well as full automation, will broaden the impact and enhance the throughput of this imaging modality further. Furthermore, the microscope is well-suited for integration into correlative light and electron microscopy (CLEM) workflows owing to the near non-destructive nature of SXT imaging, relatively small size of the microscope, and cryo-vitrified sample handling, all of which are highly compatible with cryo-CLEM workflows.

One particularly promising correlative light, electron, and X-ray microscope (CLEXM) direction for lab based soft X-ray tomography is tissue imaging. Presently, workflows including high pressure freezing followed by cryo-FIB milling and tissue lift outs or cryo-FIB milling of tissue pillars imaged by fluorescence microscopy and soft X-ray tomography followed by thinning and cryo electron tomography are being tested and fine-tuned.

Lab-based soft X-ray tomography also aligns with the growing emphasis on studying cell heterogeneity and individual cell morphology, key areas of focus for large-scale projects such as the Human Cell Atlas and the pancreatic beta cell consortium.^39,40^ Its ability to image intact, cryo-preserved cells in three dimensions without the need for fixation, staining, or sectioning makes it invaluable for studying structural variations that influence cell and tissue function, nanodrug delivery and toxicity, and disease progression. This capability is essential for understanding complex biological systems at the single-cell level, a critical aspect of modern cell biology and precision medicine.

Integrating advanced computational methods for image registration, segmentation, and analysis, such as deep learning techniques, will further expand the scope of soft X-ray tomography in biological research. This combination of SXT with machine learning has already proven fruitful for in-depth interpretation of synchrotron-generated X-ray image data of cells and will be a valuable addition to a lab-based soft X-ray tomography system in the near future.^26,41,42^

## Materials and Methods

### Yeast sample preparation

BY4741 yeast was incubated at 30°C in YPD media consisting of 2% D(+)-glucose monohydrate (Merck, 1.08342.1000), 2 % Bacto Peptone (BD Chemicals 211,677), 1 % yeast extract (Merck, 1.03753.0500), and 0.02 % adenine (Sigma-Aldrich, A-2786) for 24 hours before spinning down and resuspending the cells in PBS. Lipid droplets were labelled using a final concentration of 12 µM 4,4-difluoro-1,3,5,7,8-pentamethyl-4-bora-3a,4a-diaza-s-Indacene (BODIPY 493/503; Thermo Fisher, D3922) before cells were added to polylysine-coated Quantifoil R2/2 TEM grids (Au G200F1) for plunge freezing and imaging. The grids were plunge frozen on Leica GP2 plunge freezer using 3 second blotting times.

### *Euglena gracilis* sample preparation

*Euglena gracilis* Klebs CCAP 1224/5Z was routinely cultured in *E. gracilis* medium and Jaworski’s medium mixed in 1:1 proportions.^43^ Cultures were maintained at 15°C under a 12:12h light:dark regime. Illumination was provided by cool white LED lamps with a photon flux density of 50 μmol.m^-2^.s^-1^ at the surface of the culture vessel. For plunge freezing the cells were pipetted (4 μl) on EM grids (gold/200mesh/R2.2 either UltrAufoil or Quantifoil) in a humidified chamber (98% RH), single side back blotted (20 s) before vitrification by plunging into liquid ethane using an EM GP Plunge Freezer (Leica microsystems, Austria).

### HeLa cell sample preparation

HeLa cells were cultured in Dulbecco’s Modified Eagle Medium (DMEM) (Life Technologies, 31885023) with 10% foetal bovine serum (FBS) (Life Technologies, A5256801) on Quantifoil R2/2 gold mesh EM grids (Jena Biosciences, X-103-Au200), which were glow discharged in an argon/oxygen (75/25) atmosphere for 5 minutes. Cells were seeded at 10 × 10^4^ density and cultured overnight. Fluosphere™ 100 nm nanoparticles (Invitrogen, F8784) were added at 50 μg/μl for 4 hours to ensure uptake. Nuclei were stained with Hoechst 33342 (Sigma, 62249, 0.2 μg/ml) for 1 hour, and lysosomes with LysoTracker (Thermo Fisher, L7526, 1 µM) for 30 minutes. Cells were washed with PBS and subsequently plunge-frozen using a Leica GP2 with a 6-second blotting time to enable vitrification and avoid ice crystal formation.

### H8N8 cell sample preparation

For correlative fluorescence and SXT imaging, H8N8 mammary carcinoma cells^31^ were cultured in DMEM high glucose media (Dulbecco’s Modified Eagle Medium; Roth, 9005.1), supplemented with 10% FBS (foetal bovine serum; Merck, S0615), on Quantifoil SiO2 Au 200 R1.2/20 grids. The grids were prepared by glow discharging in an argon/oxygen atmosphere (75/25) for a period of two minutes, followed by a coating of 50 µg/ml fibronectin for 30 minutes (Merck, F1141-1MG). This was followed by a washing step with PBS to enhance cell adherence. The cells were then added to the grids and cultured overnight.

[ZrO]^2+^[(CMP)_0.99_(DUT647)_0.01_]^2-^]^2^ IOH-NPs were added to the cells at a concentration of 10 µg/ml and incubated for a period of four hours to ensure sufficient uptake. The lipid droplets were labelled using a final concentration of 5 µM 4,4-difluoro-1,3,5,7,8-pentamethyl-4-bora-3a,4a-diaza-s-indacene (BODIPY 493/503; Thermo Fisher, D3922). Nuclei were labelled using a final concentration of 5 mM Hoechst (Thermo Fisher, 62249). Both stainings were incubated for 30 minutes. For cryopreservation, plunge freezing on a Leica GP2 with a blotting time of 4 seconds was performed to achieve vitrification, preventing ice crystal formation and preserving the ultrastructure.

### Image registration using lipid droplets

The first co-registration step involves the projection of the X-ray images using a minimum intensity projection (MIP). Subsequently, a Gaussian filter is employed for smoothing, followed by CLAHE for contrast enhancement, and global Otsu thresholding to segment the lipid droplets of high X-ray absorption. The threshold is filtered to remove small or irrelevant segments, as well as regions that are connected to the image borders. For the fluorescence microscopy images, lipid droplets were stained with BODIPY and are clearly visible as green areas in the image. A region of interest (ROI) was delineated manually to focus on relevant areas. The FM images are processed similarly as the SXT stacks (Gaussian and CLAHE filtering, and conversion into a binary image using the Otsu method). To superimpose the images, the FM images are scaled to align with the X-ray data. A preliminary alignment is conducted using phase cross-correlation, followed by iterative optimisation of shifting and scaling to achieve precise overlay based on a cost function. Subsequently, the lipid droplets in both images are determined and aligned using a Radial Basis Function (RBF).

## Acknowledgments

S.O’C., D.R., A.E., P.S., W.F., T.M.E., K.F., and S.K. acknowledge funding from the European Commission for the development of the SXT 100 microscope and correlative workflows through the European Union’s Horizon 2020 Research and Innovation programme (grant agreement No. 101017116). M.C.B., J.C.S., M.S., A.W., T.M.E., K.F., and S.K. acknowledge funding from the European Union’s Horizon Europe Research and Innovation programme under the Marie Skłodowska Curie Actions Doctoral Networks (MSCA DN, agreement No. 101120151). D.W. acknowledges funding from the Danish Research Council (grant ID: 2032-00139B).

## Author contributions

S.O’C. and D.R. managed sample transfer and microscope optimisation, and all the data collection,

M.K, J.G., prepared samples, interpreted SXT and cryoFM data,

K.T. prepared samples, interpreted SXT and cryoFM data,

J.M.E. segmented yeast tomograms,

M.C.B., prepared samples, interpreted SXT and cryoFM data, wrote the manuscript,

L.F. developed an automated co-registration of florescence and soft X-ray images containing lipid bodies, performed the co-registration of the SXT and cryoFM-data

L.H., M.S., prepared samples, interpreted SXT and cryoFM data,

C.F. provided and prepared the IOH-NPs,

F.A. performed *in vitro* studies with IOH-NPs,

A.E. developed fiducial-free alignment of tomographic tilt series,

P.S., K.F., F.O’R developed the SXT-100 microscope, F.O’R also wrote the manuscript,

T.ME. managed the lab-based microscope development project,

R.F. interpreted SXT and cryoFM data, wrote the manuscript,

D.W. interpreted SXT and cryoFM data, wrote the manuscript,

J.C.S. interpreted SXT and cryoFM data, wrote the manuscript,

A.W. coordinated and provided funding for the IOH-NP project and image registration, interpreted SXT and cryoFM data, wrote the manuscript,

S.K. coordinated the imaging project, processed cryoFM images and soft X-ray tomograms, aligned visible and soft X-ray data, wrote the manuscript

