## Supplemental Information for "Demonstrating Soft X-Ray Tomography in the lab for correlative cryogenic biological imaging using X-rays and light microscopy"

### Supporting information

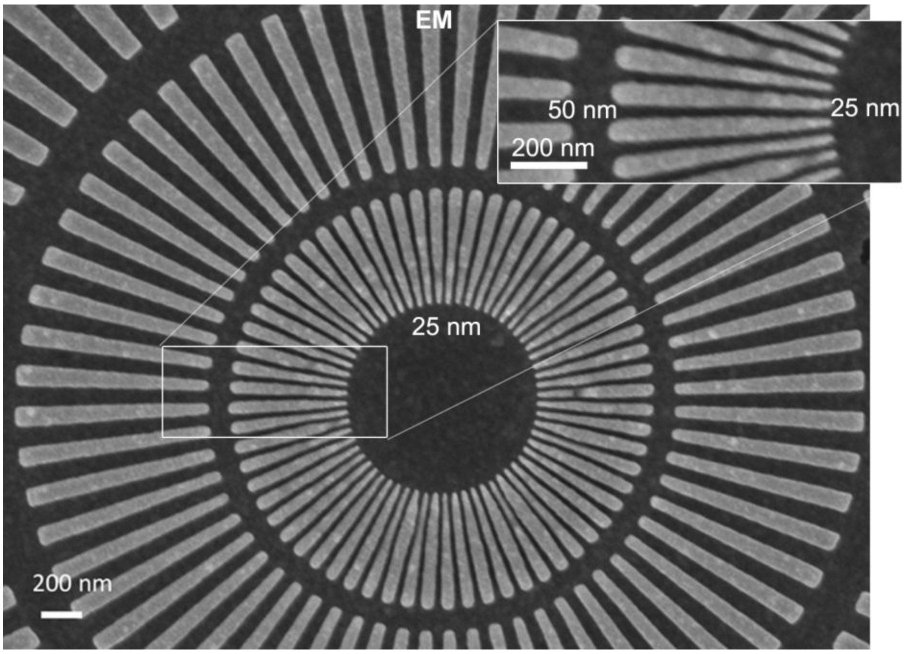


Fig. S1 A scanning electron microscope (SEM) image of the Siemens Star shown in the main text. Notably, the 50 - 25 nm ring has an uneven duty ratio of lines and spaces.
